# Waning grasslands: a quantitative temporal evaluation of the grassland habitats across human-dominated upper Gangetic Plains, north India

**DOI:** 10.1101/2021.10.10.463811

**Authors:** Shrutarshi Paul, Sohini Saha, Parag Nigam, SK Zeeshan Ali, Navendu Page, Aamer Sohel Khan, Mukesh Kumar, Bilal Habib, Dhananjai Mohan, Bivash Pandav, Samrat Mondol

**Affiliations:** Wildlife Institute of India, Dehradun, Uttarakhand; Uttar Pradesh Forest Department, Uttar Pradesh

**Keywords:** LULC changes, Temporal dynamics, Grassland-cropland mosaics, Habitat conversion, Vegetation composition, Riverine ecosystem

## Abstract

Grassland habitats currently face severe anthropogenic exploitations leading to cascading effects on the survival of grassland-dependent biodiversity globally, particularly in non-protected areas. Significant amount of such biodiversity-rich grasslands in India are found outside protected areas but lack quantitative information on their status. We evaluated the current and historical (30 years) status of the grasslands using a combination of intensive field surveys and GIS tools across one of the most fertile, human-dominated region: the upper Gangetic Plains of north India. On-ground mapping and visual classifications revealed 57% decline in grassland habitats between 1985 (418 km^2^) and 2015 (178km^2^), mostly driven by conversion to croplands (74% contribution). Radio-telemetry data from the largest endemic cervid swamp deer (n=2) showed grassland-dominated average home range (50% BBMM) size of 1.02 km^2^. The animals highly preferred these patches (average Ivlev’s index- 0.85) and showed highest temporal continuity (88%) compared to other LULC classes. Camera trapping within the core habitats suggests critical use of these patches as fawning/breeding grounds. Habitat suitability analysis indicates only ∼18% of the entire area along Ganges is suitable for swamp deer. Accurate mapping (86% accuracy) and characterization of four major grass species revealed a total 144.04 km² vegetation area, dominted by *Saccharum* sp. (35%). We recommend protection and recovery of these critical grassland patches to maintain ‘dynamic corridors’ and other appropriate management strategies involving multiple stakeholders to ensure survival of this critical ecosystem. Such evaluations, if spatially expanded, would be critical to restore this rapidly vanishing ecosystem worldwide.

## 1. Introduction

Grassland ecosystems, covering about a quarter of the earth’s landmass, play critical role in carbon sequestration, retain high biodiversity, act as valuable habitat and provide resources to the livestock population globally (Suttie et al., 2005). However, continuous anthropogenic exploitations including conversion to agriculture land, fragmentation and over-grazing have resulted in massive loss of grassland habitats during last century (White et al., 2000; Peters et al., 2006; Ceballos et al., 2010; Macdonald et al., 2013; Ripple et al., 2015; Abate and Angassa 2016). Recent reports suggest that ∼50% and ∼16% of the temperate and tropical grasslands, respectively, have been converted to agricultural lands (World Conservation Monitoring Centre, 1992), making this one of the most imperiled ecosystems globally. Such huge loss of habitats has also affected the survival of the grassland-dependent fauna (∼5% of bird species and ∼6% of mammalian biodiversity globally) (Clark, 1989; Zavaleta and Hulvey, 2004; Ceballos et al., 2010). This problem is particularly acute in regions where significant amount of grasslands and associated biodiversity exist outside the protected area network (Henwood, 1998; Mathur and Midha, 2008; Karanth et al., 2010; Harihar, 2011; Punjabi et al., 2013; Punjabi and Rao, 2017; Dorji et al., 2019; Deák et al., 2020). Implementation of required management/ conservation plans are often challenging in these regions owing to lack of appropriate information on the status, distribution and significance of the grasslands and associated fauna (Karanth et al., 2010; Kshettry et al., 2020).

With ∼24% of the total landmass, India currently retains some of the highest biodiversity rich grassland habitats (Rawat and Adhikari, 2015) within the Indian subcontinent. However, anthropogenic pressures such as habitat conversion (to agricultural land, plantations etc.) and encroachment (human settlements and grazing) pose major conservation challenges for grassland and grassland-dependent fauna across the subcontinent. In particular, the grasslands of the Terai and upper Gangetic Plains in the northern part of India have been identified as one of the biodiversity hotspots of the country (Johnsingh et al., 2004; Bashir et al., 2012; Grimmett et al., 2013). This region represents the most fertile landscape in the country and as a consequence also the highest human density (Census of India, 2011). Following the eradication of malaria in 1950s this area underwent massive habitat changes (Johnsingh et al., 2004). Currently grasslands cover only ∼2% of the Gangetic Plains as fragmented patches (Dinerstein, 2003), and majority of them are found along the basins of Sharda-Ghagra and Ganges rivers (Task Force on Grasslands and Deserts report, 2006; Paul et al., 2018, 2020) within the states of Uttarakhand and Uttar Pradesh. While most of these grassland habitats retain high biodiversity (One-horned rhino- Sale and Singh, 1987; Mishra, 1989; Kumar et al., 2002; Sinha et al., 2011; swamp deer- Qureshi et al., 2004; Tewari and Rawat, 2013a; Paul et al., 2018, 2020; Bengal florican- Rahmani, 2001; Sivakumar et al., 2014, swamp francolin- Javed et al., 1999 etc.), the two basins experience very different management regimes, wherein majority of the grassland habitats in Sharda basin are within protected areas while large parts of the grassland habitat in the Ganges basin are unprotected (Paul et al., 2020). Recent studies (Paul et al., 2018, 2020) have identified the potential grassland habitats across the landscape and mapped some of the grassland patches in the upper Gangetic basin. The study reported presence of threatened swamp deer and other fauna including hog deer (*Axis porcinus*), spotted deer (*Axis axis*), nilgai (*Boselaphus tragocamelus*), smooth-coated otter (*Lutrogale perspicillata*), fishing cat (*Prionailurus viverrinus*) and wetland birds such as sarus crane (*Antigone antigone*), black-necked stork (*Ephippiorhynchus asiaticus*), lesser adjutant (*Leptoptilosj avanicus*), Pallas’s fish eagle (*Haliaeetus leucoryphus*) and bar-headed goose (*Anser indicus*)) in these grasslands despite various forms of severe anthropogenic pressures (Paul et al., 2018, 2020). As many of these species are obligate grassland-dependent species, it is essential to understand the dynamics and composition of these grassland patches. Such information will enable us to develop effective management plans for the grassland habitats at landscape scale and eventually help in conservation of the associated faunal biodiversity.

In this paper, we used a multidisciplinary approach combining field and GIS-based tools to evaluate the current and historical (past 30 years) status of the grasslands covering both protected and unprotected areas in the upper Gangetic Plains. We conducted extensive fieldwork to map all the grassland patches along river Ganges between Haridwar (in Uttarakhand) and Garhmukteshwar (in Uttar Pradesh), as this region was identified as major swamp deer habitat earlier (Paul et al., 2018, 2020). Further, we used radio-telemetry and habitat suitability modeling to understand swamp deer habitat use patterns and identify suitable habitats in the study area. As swamp deer are obligatory grassland dwelling species, we generated a vegetation map of these grasslands by assessing four major species that are present in swamp deer habitat (Khan and Khan, 1999; Khan et al., 2003; Tewari and Rawat, 2013b). Further, we discussed the conservation implication of these results and suggested management recommendations to ensure long-term survival of the grassland habitats and associated biodiversity.

## 2. Materials and methods

### 2.1. Research permissions

All required permissions for fieldwork and sampling were provided by the Uttarakhand (1575/C-32, 978/6-32/56) and Uttar Pradesh Forest Departments (Permit Nos.; 2233/23-2-12; 3438/23-2-12). The radio-collaring permission was provided by Ministry of Environment Forest and Climate Change (MoEF&CC), Government of India and Uttarakhand Forest Department (Permit No: 1-76/2017WL).

### 2.2. Study area

This study was conducted in part of the northern swamp deer habitat along the river Ganges covering the states of Uttarakhand and Uttar Pradesh. The entire area covers ∼3173 km^2^ between Jhilmil Jheel Conservation Reserve (JJCR), Haridwar (in Uttarakhand) and Hastinapur Wildlife Sanctuary (HWLS) (covering parts of the districts Muzaffaranagar, Bijnor, Meerut, Amroha and Hapur in Uttar Pradesh). The study area includes 1677 km^2^ of protected (HWLS and JJCR) and 1496 km^2^ of non-protected area. The river Ganges flows through the centre of the entire study area (∼180 km length) and joined by it’s tributaries Banganga and Solani in Uttar Pradesh. Our earlier surveys identified that the grassland patches are restricted to 5-6 kms from both banks of the rivers (Ganges, Solani and Banganga) and this region is the only major swamp deer habitat along the Ganges (Paul et al., 2018, 2020). The study extended up to a distance of eight km from either banks of these three rivers (Supplementary Fig. A1). The study area delineated in this fashion extends from 29°79′99″N, 78°21′71″E to 28°77′34″N, 78°13′57″E. This region represents one of the most densely populated areas in the entire country (population density of 1164 people/ km^2^ compared to national average of 382 people/ km^2^ (Census of India, 2011). Agricultural fields, villages, townships, grassland patches, few scattered scrublands and forests make up majority of the mosaic in this human-dominated region.

### 2.3. Grassland status: Land-use Land-cover classification

Significant numbers of the existing grasslands habitats are outside the protected area boundaries where they have been subjected to heavy anthropogenic exploitations over the last few decades (Paul et al., 2018, 2020). Initial assessment through Google Earth imageries revealed that the study area broadly comprises of following land-use patterns-cropland, grassland, waterbody, settlement, forest and scrubland. Subsequently, the two banks of the Ganges (river stretch of 180 km) and its tributaries Banganga and Solani (rivers stretches of 66 km and 70 km) were extensively surveyed on foot, tractor or boat and coordinates of different LULC classes were recorded as ground points. A total of 3253 km of survey spanning the entire study area was conducted between 2015-2016 for full spatial coverage. A maximum distance of 8 km was surveyed on both sides of Ganges and its tributaries. We collected a total of 656 GPS points for reference, representing six different LULC classes mentioned above.

Further, we downloaded Landsat images (USGS) for 2015 (Landsat 8) with 30 m resolution. After radiometric correction of images for the focused study area, we visually interpreted the Landsat images using spectral characteristics (size, tone, texture) of different LULC classes and field knowledge (656 GPS points) (Mas and Ramírez, 1996; Babu et al., 1997; Schroeder et al., 2006; Al Rawashdeh, 2012). Such visual classification was opted, as extensive field information was available to us, where the classification was done at a scale of 1:50000 (Puig et al., 2002; Al Rawashdeh, 2012) and six above-mentioned LULC classes were categorized. Once the classification was done, we randomly generated another 100 points representing different LULC classes in ArcGIS 10.2.2 and further validated them through additional field surveys. Accuracy of classification was defined as:

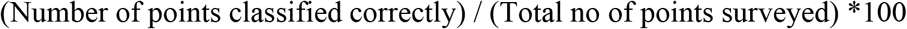

The total area of different LULC classes was calculated on a GIS domain (ArcGIS 10.2.2) and percentage cover of each class was represented through Pie charts.

Similarly, the LULC maps of 2005 (Landsat 7), 1995 and 1985 (Landsat 4-5 TM) were generated through visual interpretation using 2015 classification as reference. We calculated the net changes in each LULC class in the last 30 years in a GIS domain. As majority of changes in the landscape were restricted to grasslands and croplands, we focused our analysis on fine-scale changes incurred by croplands and grasslands in this landscape. Analysis of interchange of classes over the years was performed using the Union tool in ArcGIS 10.2.2.

### 2.4. Assessment of grassland usage patterns by swamp deer

Swamp deer is the largest and obligate grassland dependent mammal in this area, and its future survival will entirely depend on the management of these existing grassland patches (Paul et al., 2018, 2020). To understand the importance of these habitats, we analysed habitat suitability (through MaxEnt modeling) and assessed the habitat use pattern of collared swamp deer over a period of around one year.

#### 2.4.1. Radio-telemetry: Capture and tracking

Standard approaches of radio-collaring (darting from vehicle, elephant back, hides etc.) are extremely challenging for swamp deer due to their open habitat preferences (Qureshi et al., 2004.) and sensitivity to human presence. We used the drive-net method to capture and collar two female swamp deer. This approach has been used successfully in similar species globally (White-tailed deer-Locke et al., 2004; Argali-Ramey II et al., 2004; Saiga antelope- Berger et al., 2010). The collaring operations were conducted in JJCR, Uttarakhand during May-June 2018. The capturing of the individuals was performed using about 150 m long nylon-mesh drive nets. The animals were acclimatised to the nets for about three months before final capturing operations were conducted. A group of ∼60 field staffs were used in the driving process. Once captured, the animals were blindfolded and administered with a mild dose of sedative Azaperone (40mg/ml dose) and fitted with GPS Vertex Plus satellite collars (Vectronic Aerospace). The collars were set to provide information on latitude, longitude, time and temperature at every 2-hour intervals. The two collared animals were monitored for 14 months (Female 1) and 11 months (Female 2), respectively, using both on-ground tracking as well as the satellite information.

#### 2.4.2. Home range estimation

Using the Brownian Bridge Movement Model (BBMM), we estimated home ranges of two collared swamp deer during the entire period of their data transmission (May 2018-July 2019). The BBMM requires (1) sequential location data (2) estimated error associated with location data and (3) grid cell size assigned for the output utilization distribution. We selected a fine-scale grid cell size of 30m for the analysis as we did not have any prior information on swamp deer movement patterns. We prepared 50% and 95% BBMM to represent the core area of use and the standard home range size using BBMM package (http://cran.opensourceresources.org) in R software (Horne et al., 2007; Fischer et al., 2013). Further, we plotted these home ranges within different LULC classes of the study area.

#### 2.4.3. Habitat selection patterns at spatial and temporal scales

We assessed swamp deer habitat selection using Ivlev’s electivity index (Ivlev, 1961; Jhala et al., 2009). The Ivlev’s index is calculated as:

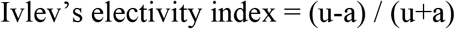

where, u= proportion of GPS points in a particular LULC class (use); a= proportion of a particular LULC class available (availability). The electivity value ranges from −1 to +1 whereas −1 signifies habitat avoidance and +1 indicates exclusive use, while 0 represents habitat used in proportion to availability (Ivlev, 1961; Kong et al., 2018). This process can be defined at the level of landscape/study area (First-order selection), the home range of an individual animal or group (Second-order selection) and usage of specific components within the home range (Third-order selection) (Johnson, 1980; Krausman, 1999; Buskirk and Millspaugh, 2003). For initial analyses, we plotted the proportion of all GPS locations from both collared individuals in each LULC class to get a qualitative idea of habitat selection without considering habitat availability. However, for more quantitative analyses of first and second order selections using Ivlev’s index, we defined habitat availability at landscape (First-order selection in the entire study area) or local (Second-order selection within 95% and 50% BBMM home ranges) scales. We also plotted the proportional use versus availability of different LULC classes (grassland, cropland and waterbodies) at the three scales (landscape, 95% BBMM and 50% BBMM).

To understand swamp deer temporal habitat use in different LULC classes we conducted a temporal trajectory path continuity analysis (Fuller and Harrison, 2010; Parent et al., 2013; Qi and Du, 2013). We extracted the data locations of collared individuals covering each LULC class and used trajectory path tool in ArcMet package (ArcGIS 10.2.2) to create trajectory movement paths and evaluated the proportion of points that could not be joined temporally (isolated points) in each habitat class (Wall et al., 2014).

Finally, we deployed camera traps to understand human-swamp deer temporal overlaps at a fine scale in the non-protected grassland patches. These grassland patches were selected based on the GPS locations of the collared animals. In order to increase photo capture rate, the infrared motion-sensor cameras (Cuddeback) were strategically placed in accessible areas covering grassland trails, grassland-cropland interphases and areas between two small grassland fragments. The camera-trapping was conducted in three sessions between July 2018 and May 2019 with a total effort of 376 trap nights (Session 1 in July: 07 cameras, 84 trap nights, Session 2 in November: 11 cameras, 148 trap nights; Session 3 in April: 12 cameras, 144 trap nights). We analysed the data using Oriana software v2.0 (Gómez-Ortiz et al., 2019) and calculated the temporal overlap between human and swamp deer using CamTrap package implemented in R software (Niedballa et al., 2016) (parameters; deltatime>15mins, overlap estimator used=Dhat4, 95% CI of overlap generated by bootstrapping 999 samples) (Niedballa et al., 2016; Dorning and Harris, 2019; Allen et al., 2021; Vilella et al., 2021). Significance in temporal activity difference between swamp deer and human was assessed using Watson U^2^ test (Rubio et al., 2017).

#### 2.4.4. Habitat Suitability

We conducted fine-scale habitat suitability analyses at the landscape level as our inferences regarding swamp deer home ranges and habitat use are based on insufficient sample size (n=two collared individuals). For this, we also used earlier published data on swamp deer presence points (direct sighting, antlers and genetically identified pellets) (Paul et al., 2018, 2020). The points were spatially filtered and modeled in Maxent software (1×1 km^2^ grids) (Phillips et al., 2006) using 6 covariates viz. six LULC classes, Elevation (m), Annual Precipitation (mm), Human Population density (per km^2^), Nightlight and Distance from rivers (km) after checking for autocorrelation between variables. The final performance of the model was assessed using AUC curves. The parameter and model settings were used following Paul et al. 2020. We pooled all the prediction probabilities (based on 10 percentile training presence logistic threshold value) to create areas with high and low suitability based on the mid-value of prediction range (Morales et al., 2017; Paul et al., 2020). Further we assessed the total area of suitable habitat left for swamp deer both inside and outside protected areas and identified the critical areas which need management attention.

### 2.5. Vegetation composition of grasslands

The fragmented grassland patches are the only available resources to the swamp deer population in this region. Therefore, apart from their distribution and status it is also important to characterize the vegetation compositions of these patches due to the obligatory species dependence on them. Earlier studies indicated a number of grass species (*Phragmites* sp., *Typha* sp., *Saccharum spontaneum*, *Saccharum bengalense* etc.) within the Gangetic grassland habitats with swamp deer presence (Tewari and Rawat, 2013a; Paul et al., 2020). Our initial assessment based on reflectance values of the Google earth imageries indicated potentially different but unclear patterns of species homogeneity/heterogeneity. We used a combination of field sampling and vegetation indices to generate vegetation map and assess the species composition of these grasslands (Xie et al., 2008; Kassa et al., 2016).

The vegetation sampling was conducted in two phases. First, we surveyed all the grassland patches between Haridwar and Bijnor barrage in the upper Gangetic Plains, where 98 (5X5 m²) training vegetation plots were laid following a stratified random framework. The numbers of plots were proportionate to vegetation heterogeneity of the patches. Within each plot, individual numbers of stems/tufts of *Phragmites* sp., *Typha* sp., *S. spontaneum*, *S. bengalense* were counted. In homogeneous patches 1×1 m^2^ plots were laid at each corner of 5X5 m^2^ plots and the species were counted and later extrapolated. Subsequently, each plot was assigned as one of the following four vegetation classes (based on a threshold of 80% coverage) (Wikum and Shanholtzer, 1978):

1. Pure *Phragmites*/ *Typha* (consists of pure *Phragmites* sp. and/or *Typha* sp.)
2. Pure *Saccharum* (consists of pure *S. bengalense* and/or *S. spontaneum*)
3. *Phragmites* Dominated Mix (dominated by *Phragmites* sp and/or *Typha* sp. with presence of *S. bengalense* and/or *S. spontaneum*)
4. *Saccharum* Dominated Mix (dominated by *S. bengalense* and/or *S. spontaneum* along with *Phragmites* sp. */ Typha* sp.)

We used Normalized Difference Vegetation Index (NDVI) to generate imagery based vegetation map for this region (Bannari et al., 1995; Eastman et al., 2013; El-Gammal et al., 2014). We downloaded Landsat images (Landsat 8-USGS, 2018) with 30 m resolution and masked only the grassland areas from obtained NDVI image. Further, we trained NDVI with the GPS points of different field-collected vegetation data and reclassified NDVI values for the entire study area into above-mentioned four grassland types. Subsequently, we conducted the second phase of vegetation sampling to assess the accuracy of our NDVI classification. This sampling was focused from the south of Bijnor Barrage to Garhmukteshwar, Uttar Pradesh (the end of our study area), where we laid 111 plots and assigned them into one of the four vegetation classes mentioned earlier. The overall accuracy of our vegetation classification was defined as

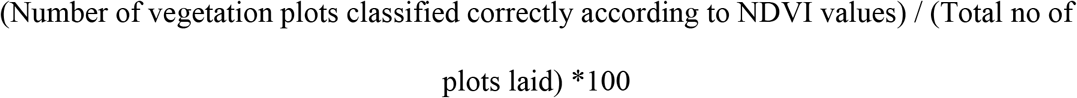

Finally, we performed a multivariate ordination analysis to detect difference in species composition among four vegetation classes and estimated relative abundance of the major grass species in the vegetation plots. We performed NMDS analysis using Scatterplot3d and Vegan package in R (Clake, 1993; Cavender et al., 2014; Oksanen et al., 2020; https://CRAN.R-project.org/package=vegan) and tested the significance in difference of vegetation between the groups using bootstrapping PERMANOVA test (Anderson, 2014; https://CRAN.R-project.org/package=PERMANOVA). For relative abundance analyses, we ran Box Plot in R (Kua et al., 2020). Further, Kruskal-Wallis test was performed to indicate the significance in differences of mean relative abundance values between four species of each class (Kruskal, 1964; Reinecke et al., 2013). Additionally, we calculated the area coverage of different types of vegetation classes in the study area using ArcGIS 10.2.2. We also calculated Ivlev’s index (1^st^ and 2^nd^ order selections) for preference of any specific vegetation type within the grasslands in the summer season (as we only had ground validated NDVI based vegetation maps of April and May representing summer).

## 3. Results

### 3.1. Land-use land-cover change at temporal scale (1985-2015)

Our 656 field-collected data points matched the different LULC classes (cropland, grassland, forest, waterbody, settlement and scrubland) in Google Earth, and helped us to visually classify the Landsat image. The second survey to test the accuracy of LULC classification with 100 randomly selected points resulted in 91% success. In 2015, cropland was the most dominant LULC class (76%), followed by waterbodies (7%), forest (6%), grassland (6%), settlements (4%) and scrublands (1%), respectively. The LULC change detection analyses showed an increase in cropland cover from 69% (1985) to 76% (2015), whereas the grassland cover reduced from 13% (1985) to 5% (2015) (Fig. 1). As the other habitat classes did not show much temporal changes, they were not considered for further analyses.

**Fig. 1:**
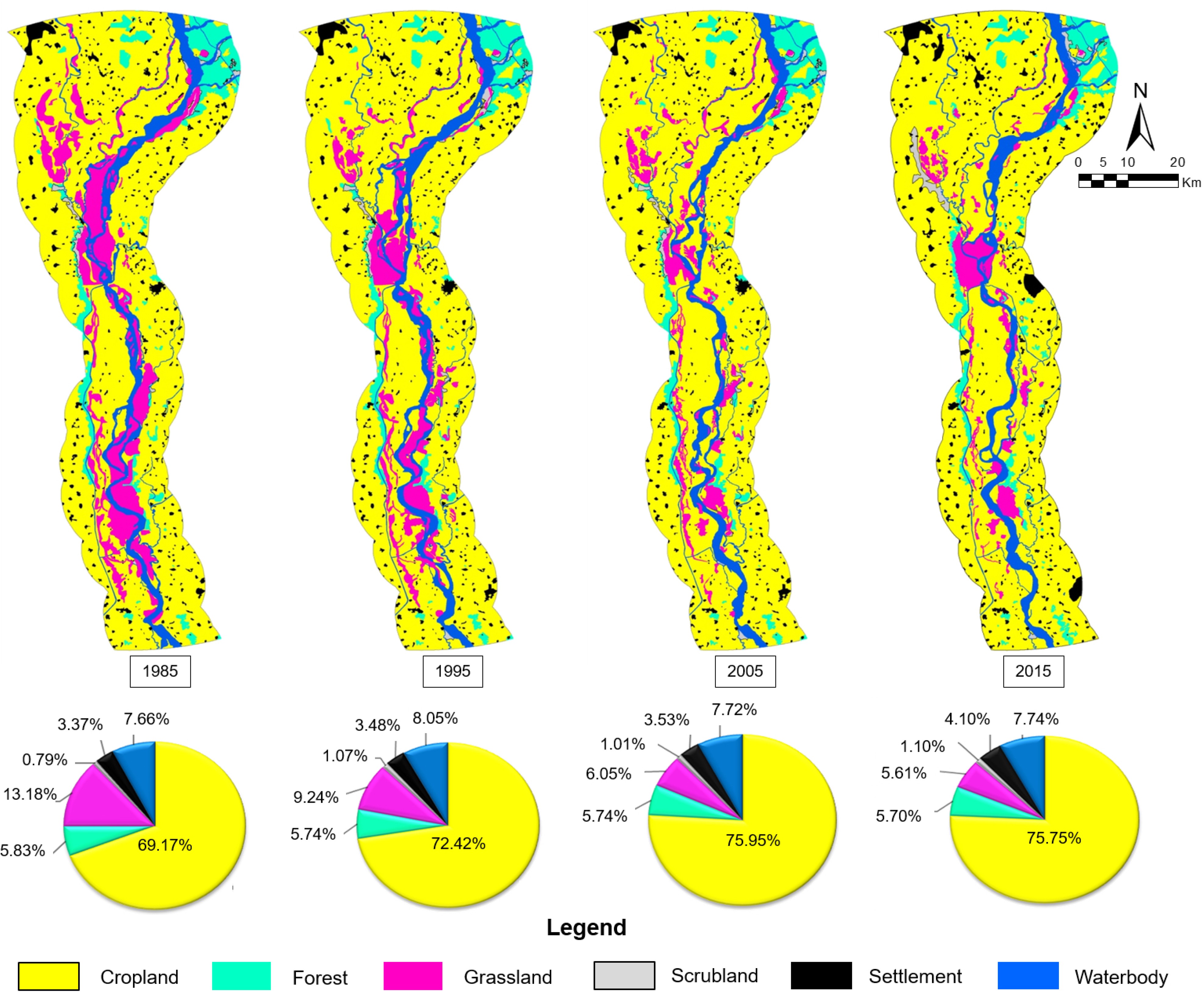
Changes in the landuse-landcover patterns in the study area over three decades (1985-2015). Below pie-charts represent the percentages of different LULC classes for respective decades, indicating major reduction in grassland habitats.

Further, fine-scale assessment of the grassland-cropland changes within the protected (∼1636 km^2^ area within HWLS covering 52% of total landscape) and non-protected (∼1496 km^2^ covering 47% of landscape) areas showed that ∼193 km^2^ and ∼47 km^2^ of grassland area was lost respectively, during 1985-2015. Jhilmil Jheel Conservation Reserve (JJCR) (∼41 km^2^ area covering 1% of the landscape) was not considered in this analysis as it was designated as a protected area in 2005 (Sinha and Chandola, 2006). Croplands showed a corresponding increase of ∼173 km^2^ and ∼34 km^2^ within and outside protected areas, respectively. Majority of this grassland loss was attributed to conversion into croplands (74%) followed by waterbodies (20%) and other classes (6%), whereas the increase in cropland was due to grassland loss (65%) followed by waterbodies (26%) and other classes (9%) (Table 1).

**Table 1:**
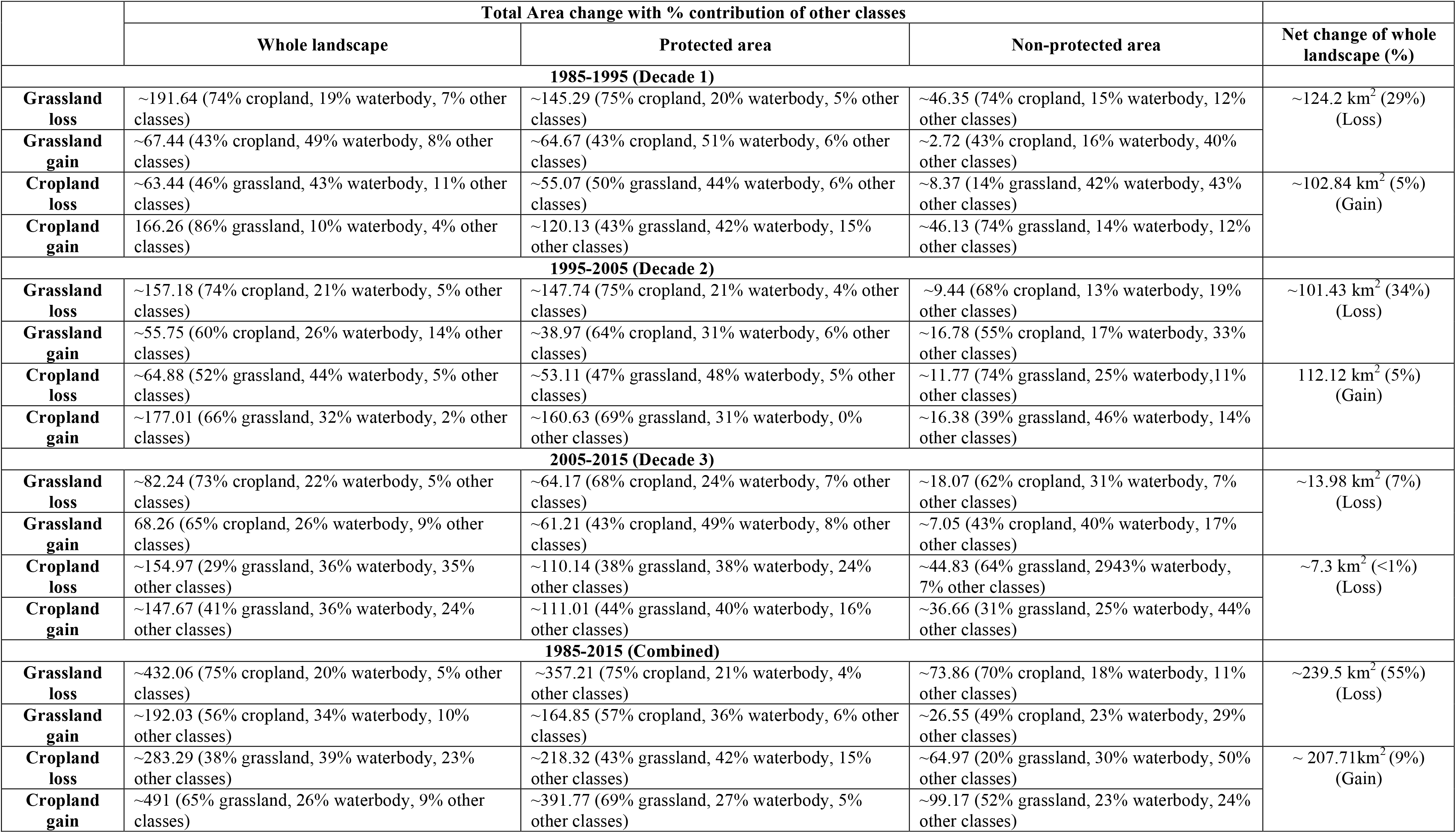
Quantification of the total changes in grassland and cropland areas and the contributing major LULC classes (grassland/cropland/waterbody) towards the changes during last three decades (1985-2015). Decade-wise and overall, changes are presented within: whole landscape, protected area (HWLS only) and non-protected areas.

### 3.2 Swamp deer ranging pattern, habitat use and suitability

Two collared swamp deer females produced 4998 (Female 1- May 2018-July 2019) and 3792 (Female 2- June 2018-May 2019) GPS locations, respectively. The average 95% BBMM home range of both individuals was 10.27 km^2^ (Female 1-8.22 km^2^, Female 2-12.33 km^2^, respectively) covering grassland (32%), cropland (30%) and waterbody (25%) habitats. However, the average 50% BBMM home range (intensive use area) was 1.02 km^2^ (Female 1- 0.58 km^2^ and Female 2-1.47km^2^, respectively) (Fig. 2A, Table 2) consisting 62% grassland, 29% cropland and 1% waterbody, indicating intensive use of the small grassland patches as core habitats.

**Fig. 2:**
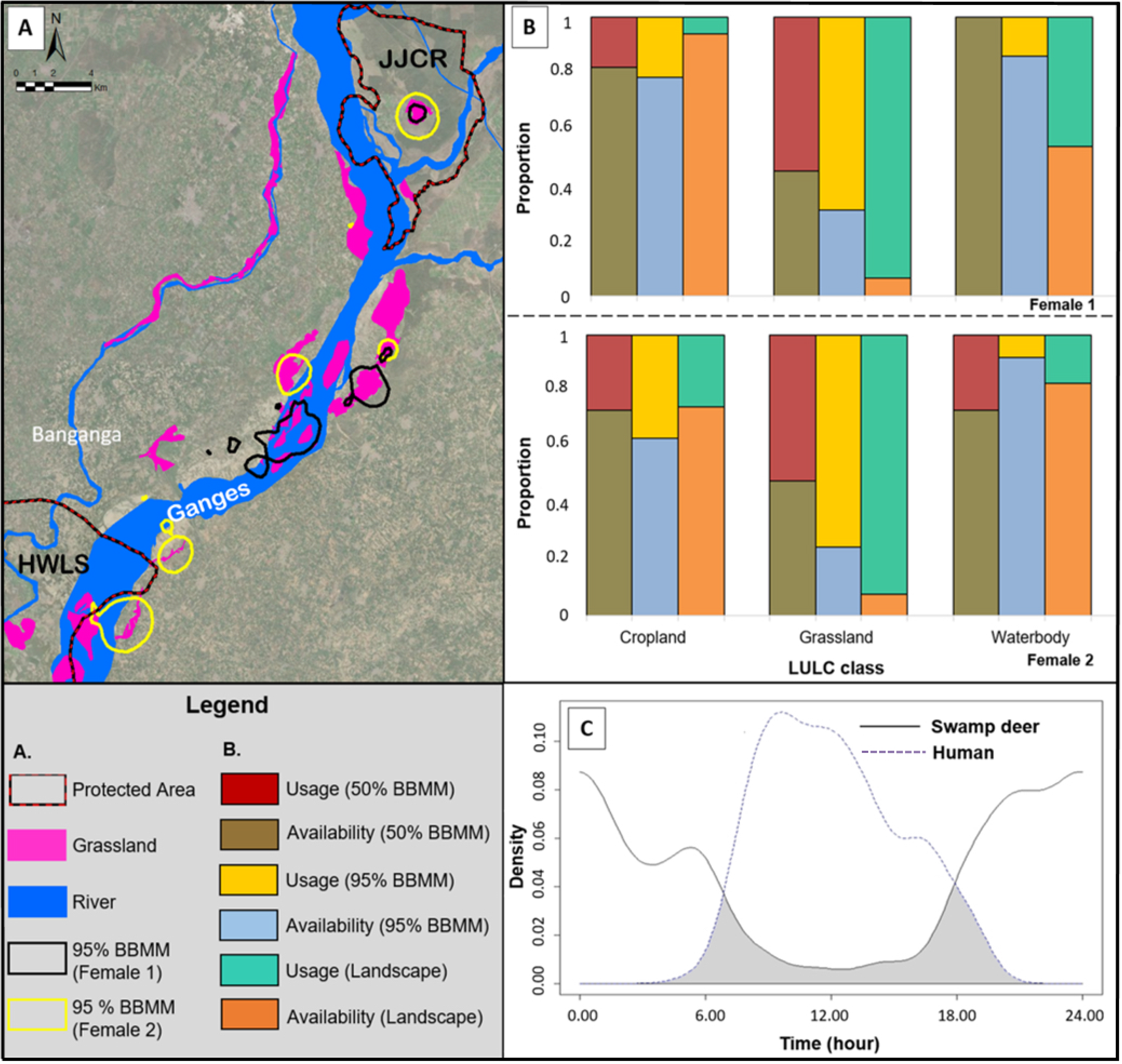
Swamp deer home range and habitat use patterns ascertained through radio-telemetry (n=2 females) (A) Spatial representation of two female swamp deer home ranges (95% BBMM), indicating use of the fragmented grassland patches in this landscape; (B) Proportion of use vs. availability of grassland, waterbody and cropland classes for two collared females at different scales (landscape, 50% and 95% BBMM). The results suggest higher use of grasslands against their availability; (C) Temporal segregation between swamp deer and human suggesting avoidance to human presence by the species.

**Table 2:**
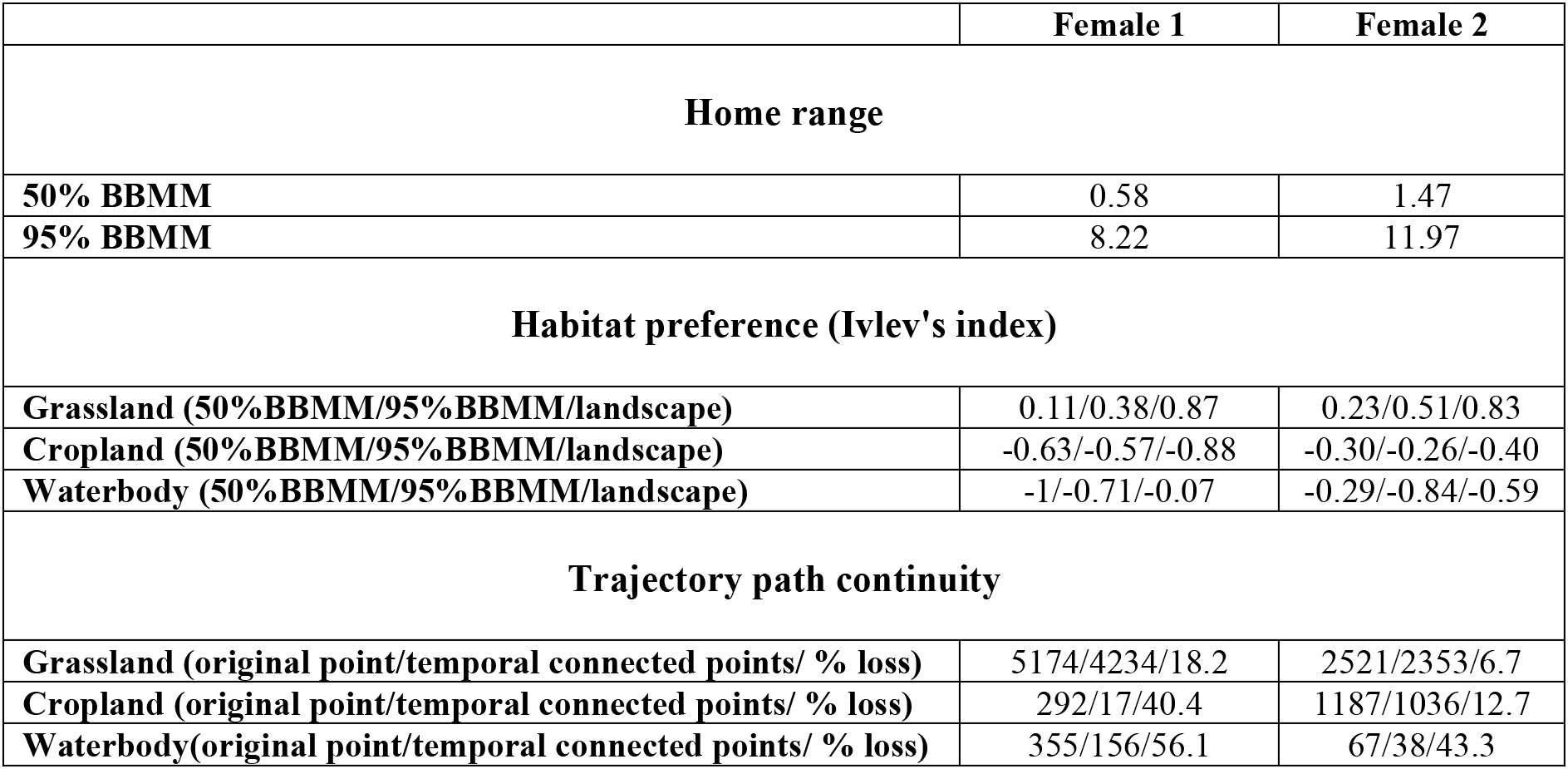
Details of swamp deer (n=2 females) home ranges (50% and 95% BBMM), habitat preferences (Ivlev’s Index) and temporal continuity of trajectory paths. The habitat preferences have been calculated at three scales (entire landscape, 50% and 95% BBMM) and trajectory path analyses was conducted for major LULC classes (grassland, cropland and waterbody).

Qualitative analyis for habitat selection (not considering availablity) for both the collared individuals suggested 81% of GPS locations were within grassland habitats followed by cropland (15%) and waterbodies (4%). Ivlev’s index analysis (considering avialability) indicated that both animals preferred grassland habitats and avoided forests, scrublands and waterbodies. Results also incidated sporadic use of croplands. At landscape level (1^st^ order selection) the Ivlev’s index values for grassland were highest for both individuals (Female 1- 0.87 and female 2-0.83, respectively), when compared to 95% BBMM (Female 1-0.38 and female 2-0.38, respectively) and 50% BBMM Female 1-0.11 and female 2-0.25, respectively) (both 2^nd^ order selections) (Table 2). Similar trends were observed for the proportion of use versus availablity analyses (Fig. 2B).

The temporal trajectory path continuity analysis revealed that grasslands had the highest temporal continuity followed by croplands and waterbodies. For Female 1, 82% of the locations within grasslands was temporally connected followed by croplands (60%) and waterbodies (44%). Similar pattern was observed for Female 2 (Temporal continuity: 93%- grassland, 87% cropland and 57% waterbodies) (Table 2). The camera trapping exercise (effort of 376 trap nights) around the core habitats of two collared individuals resulted in 403 and 157 photo captures of swamp deer and human, respectively. The data indicated presence of small groups (3-4 individuals/group) within both the collared individuals’ core habitat regions, which were also important rutting and fawning grounds (Supplementary Fig. A2 B and C). Swamp deer activities were temporally separated (Frequency-0.94, 6:00 pm to 6:00 am) from human (Frequency-0.95, 7:00 am and 4:00 pm) (U^2^= 7.41 p<0.001) with a 21% overlap (Dhat4= 0.21, 95% CI- 0.16, 0.26) (Fig. 2C).

MaxEnt analyses indicated that most of the suitable swamp deer habitats are along the river Ganges and its tributaries, Solani and Banganga. Based on the model predictions (AUC value- 0.909) (Supplementary Fig. A3 G), only ∼18% of the entire study area was found to be suitable as swamp deer habitat. The highly suitable areas (prediction probablility between 0.56-0.95) were ∼91 km^2^ (∼3%) region, mainly between JJCR and Bijnor Barrage along Ganges, whereas the low suitable area (0.18-0.56) was ∼472 km^2^ (∼14%) stretch around river Solani and south of Bijnor Barrage (Fig. 3). Fine-scale analyses revealed that ∼27 km^2^ and ∼64 km^2^ of highly suitable habitats are present in protected and non-protected areas, respectively. Distance to water (24.9%) and grassland cover (24.7%) were the most critical predictors of swamp deer habitat suitability (Supplementary Fig. A3 A-F and H).

**Fig. 3:**
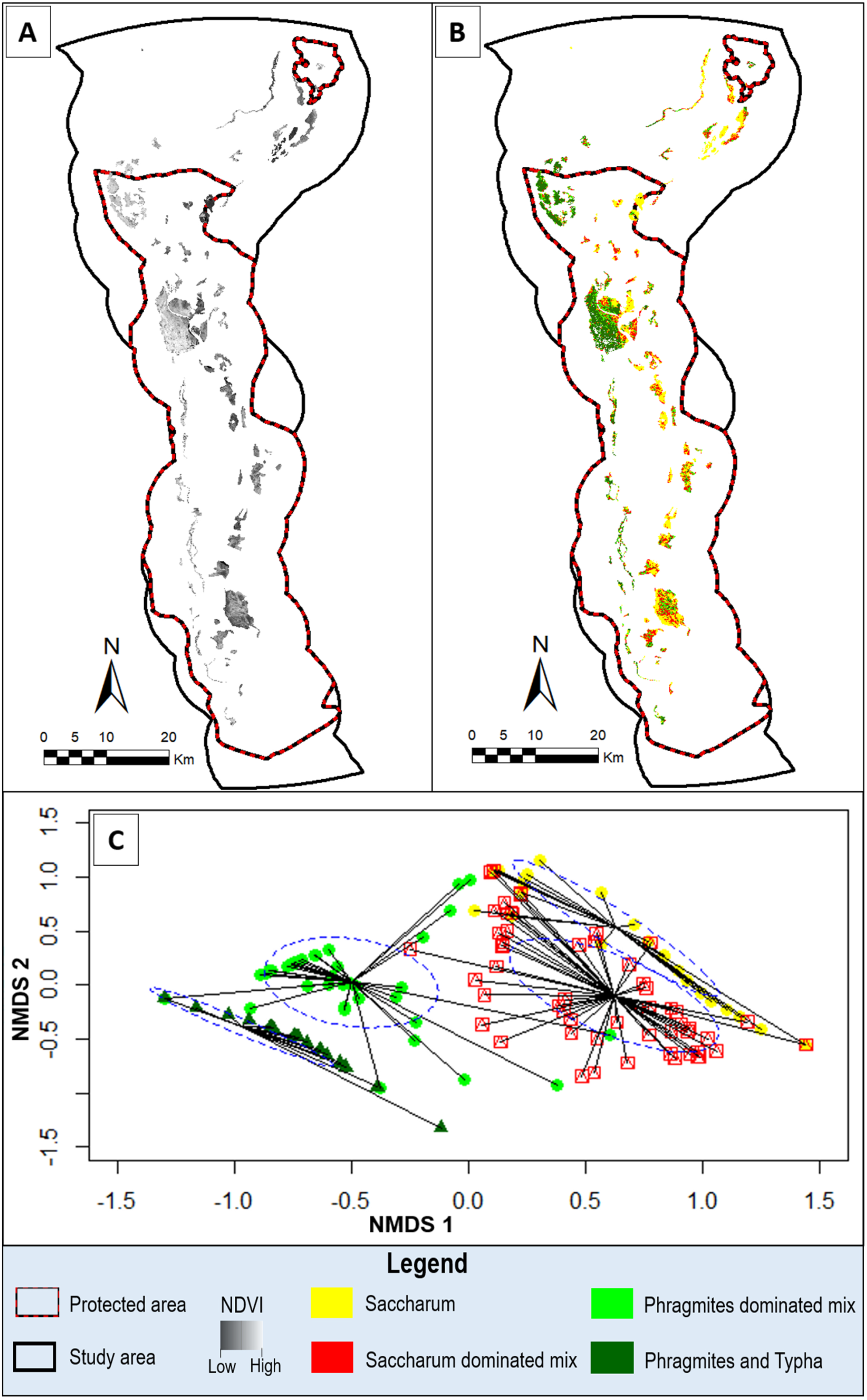
Habitat suitability map for swamp deer representing both high and low suitability areas in the upper Gangetic plains. The map suggests that significant portion of the highly suitable habitats are outside protected areas (JJCR and HWLS).

### 3.3. Species composition of the grasslands

Based on the field-collected vegetation data (n=209 plots) and distinct NDVI values, we classified the vegetation classes having following NDVI ranges: Saccharum (NDVI-0.12-0.315), Saccharum Dominated Mix (NDVI-0.316-0.39), Phragmites Dominated Mix (NDVI-0.40-0.44) and Phragmites-Typha (NDVI-0.45-0.56) (Fig. 4A and B). However, *Phragmites* sp. and *Typha* sp. showed similar reflectance values and could not be separately classified. Our second phase vegetation sampling (n=111 plots) showed 86.4% accuracy in NDVI classification.

**Fig. 4:**
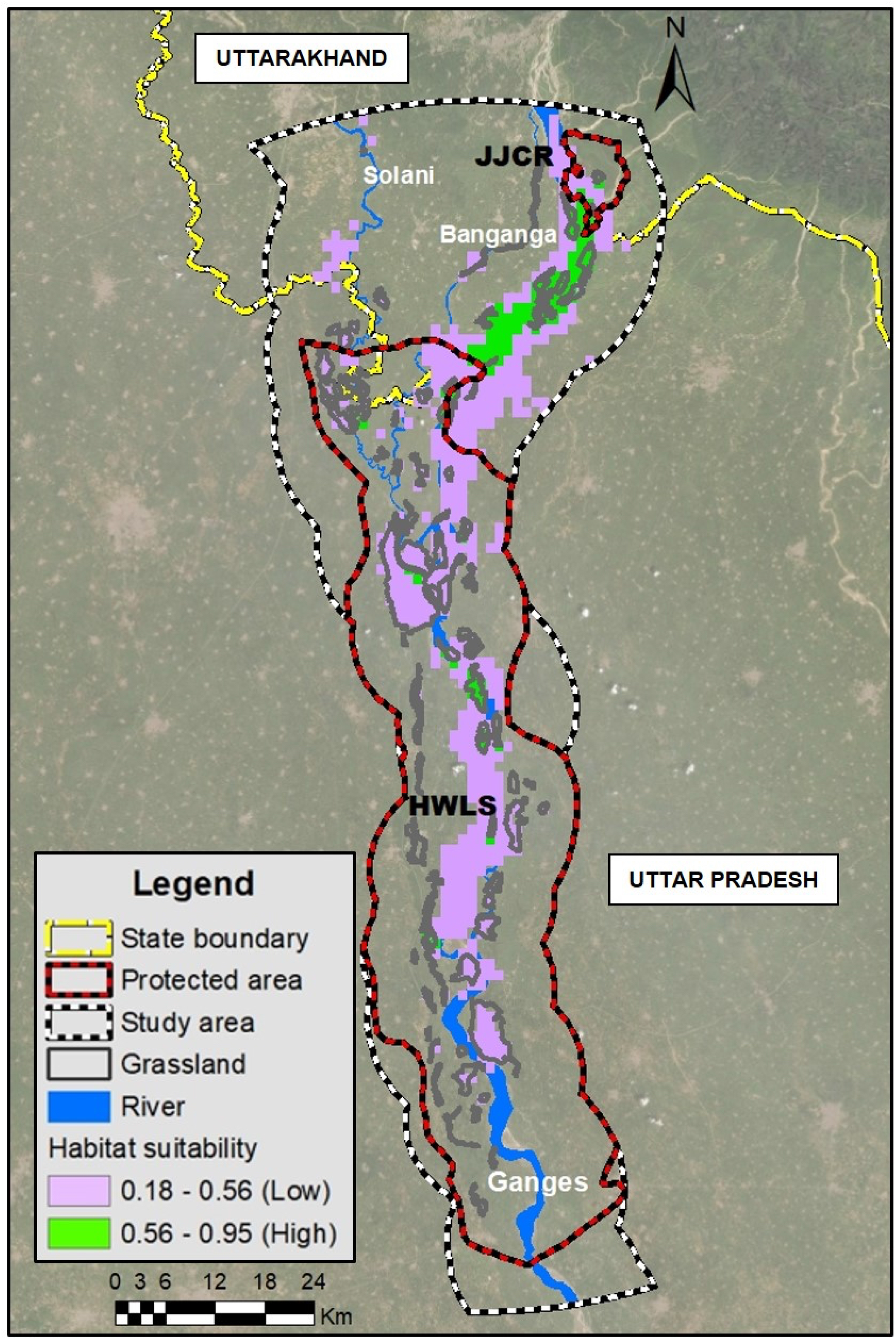
Vegetation mapping and composition analysis of four vegetation species ((Pure Phragmites/ Typha, pure Saccharum, Phragmites Dominated Mix and Saccharum Dominated Mix) in the study area. (A) Raw NDVI value-based grassland vegetation map of the study area; (B) Vegetation map with reclassified NDVI values for four vegetation classes; (C) Two-dimensional NMDS plot showing the comparison among the four vegetation types. The clusters show minimal overlap indicating discernable differences in species composition (*S. spontaneum*, *S. bengalense*, *Phragmites* sp. and *Typha* sp.) among them.

NMDS ordination (Stress value 0.13), showed three separate vegetation clusters with overlaps between Saccharum and Saccharum Dominated Mix indicating varying differences in species composition among the four classes (PERMANOVA p<.01) (Fig. 4C). Within Saccharum class, *S. spontaneum* was the most abundant among all the four species. Saccharum Dominated Mix and Phragmites Dominated Mix classes showed presence of all four species though *S. spontaneum* and *Phragmites* sp. were more abundant in their respective plots. In Phragmites-Typha class, most abundant species was *Phragmites* sp. followed by *Typha* sp. and *Saccharum* spp. (Supplementary Fig. A4). Comparison between mean relative abundance of each species within the four vegetation types also revealed similar patterns (Supplementary Table A1). Total area covered by these four vegetation classes was 144.04 km². Pure Saccharum was the dominant vegetation type covering 35% of the landscape, followed by Phragmites-Typha (27%), Saccharum Dominated Mix (23%) and Phragmites Dominated Mix (15%). Based on frequency of the positive Ivlev’s index (preference) across the three levels of selection (Landscape, 95% BBMM and 50% BBMM), it was found that swamp deer (both individuals combined) habitat selection had 42% occurrence of Saccharum dominated mix followed by Phragmites dominated mix (33%), Phragmites-Typha (17%) and Pure Saccharum (8%) (Supplementary Fig. A5).

## 4. Discussion

Our study presents the most exhaustive assessment of the biodiversity-rich grassland habitats in the upper Gangetic Plains of north India. Further, our analyses revealed that in 1985 a total of ∼418km^2^ grassland area was available and the region has observed ∼57% (∼178km^2^ grassland in 2015) decline in grassland habitats during last three decades (1985-2015), mostly driven by habitat conversion to croplands (accounting for ∼74% of grassland loss). Multidisciplinary approaches involving field surveys, radio-collaring, camera trapping and habitat suitability analysis revealed that the existing fragmented grassland patches are highly preferred and critical breeding/fawning grounds for swamp deer (Supplementary Fig. A2 B and C). Given that currently only ∼5% of the study area is grassland habitat, these patches are absolutely essential for future swamp deer survival. Finally, we developed a vegetation map with 86.4% accuracy focusing on four major species known to characterize swamp deer habitats and pure Saccharum was the dominant vegetation type in the study area.

Some of the critical components of this study such as habitat mapping and change detection was dependent on accurate LULC classification. We decided to use visual interpretation of Landsat imageries as this approach has been reported to show similar quality as digital classification for analysis of medium resolution satellite data, particularly when the number of LULC categories are less (Puig et al., 2002; Ghorbani and Pakravan, 2013) and detailed information on the study area are available (Puig et al., 2002). Our study landscape had only six major LULC categories and our intensive survey efforts generated fine-scale georeferenced locations of these categories that provided best quality LULC classifications for downstream analyses. However, it is important to point out that during our analyses some habitats types were merged within the six classes (for example, orchards as cropland, plantations as forest, river sandbars as waterbody etc.), as these classes could not be differentiated in the historical images. Future efforts may consider more detailed LULC classification for fine-scale analyses. Decade-wise comparative analyses of net habitat loss showed that the highest loss of grasslands occurred during 1985-1995 (∼125 km^2^), followed by 1995-2005 (∼102 km^2^) and 2005-2015 (∼14 km^2^) (Table 1). However, when observed at percentage-wise habitat loss, 1995-2005 showed the highest loss (34% loss in grasslands from ∼293km^2^ to ∼191km^2^), closely followed by 1985-1995 (29% loss from ∼418 km^2^ to ∼293 km^2^) (Table 1). Earlier reports also corroborate this pattern of less reduction in grassland habitats during last decade (Tsarouchi et al., 2014), possibly due to more frequent shifting in river courses and better management strategies (Sinha and Chandola, 2006; Bhargava, 2007.). Although we were able to compare the habitat loss for the last 30 years period, available information suggest a long history of such events in this region (Johnsingh et al., 2004). Till 1940s-1950s, this entire region was dominated with swampy grassland habitat and very high incidences of malaria infestation, resulting in rather low human density. Following the introduction of DDT (as anti-mosquito agent) and other advanced drugs post world war II, and major resettlement of people from erstwhile East Pakistan by Government of India (Johnsingh et al., 2004) this region faced a major population boom. Continuous encroachment of lands for settlement and agricultural requirements has led to severe loss of the swampy grassland habitat. Interestingly, we observed that this recent loss has also affected the protected areas of this landscape. For example, our analyses indicate an alarming loss of 193 km^2^ grassland habitat in last three decades within the boundary of Hastinapur Wildlife Sanctuary (HWLS) (representing ∼80% of total loss) (Table 1). HWLS is the largest protected area in the entire landscape and was exclusively created to protect the swamp deer (WII-EIA Technical Report, 2009) and other Gangetic fauna. Such loss in HWLS has long-term negative impact in conservation of Gangetic grassland biodiversity and thus require urgent and immediate attention.

Despite very intensive anthropogenic pressures our results show that about 13% of the grassland patches are still surviving outside protected area between JJCR and HWLS. We believe that the extremely dynamic nature of the Gangetic riverine system supports continuous regeneration of newer grassland habitats and counter the loss from cropland conversion. Earlier studies have reported how such dynamics shape the structure of riverine systems and associated human-modified LULC classes (Hazarika et al., 2015; Rai et al., 2018). Even in this study landscape, we observed that the net loss of grasslands comes from a combination of fine-scale patterns of habitat loss and gain resulting from river flow dynamics. To explain this, we have conducted a time series analysis of grassland change dynamics of a representative river island region within our landscape (Supplementary Fig. A6). This area, called Rauli Ghat (15.52 km^2^-2015) is located ∼9km upstream of Bijnor Barrage embedded between the Solani and the Ganges. A detailed, comprehensive analyses of Google Earth imagery of this area between 2005 and 2020 (four time periods: 2005-7.9 km^2^, 2010-8.71 km^2^, 2015-15.52 km^2^ and 2020-5.91 km^2^) revealed continuous change in grassland habitat area, possibly resulting from a combination of Ganges river course changes (between 2005-2015) and conversion to croplands (between 2015-2020). Such situation is very common in this entire landscape and possibly the only reason why the grassland habitats and associated fauna continue to survive under such high anthropogenic pressures. Another potential area of concern for the grasslands are certain management decisions such as plantations as part of the existing forestry practices. In many areas of this landscape, several species such as *Eucalyptus* spp., *Terminalia arjuna* (arjun), *Sengegalia catechu* (khair), *Vachellia nilotica* (babul) etc. have been planted within the core grassland patches (Holloway, 1972; Birdlife International report on Indo-Gangetic grasslands-https://www.birdlife.org/action/science/asiastrategy/pdf_downloads/ grasslandsGO2.pdf).While such plantation-based management practices can reduce encroachment threats, they alter habitat quality, soil properties, vegetation composition and affect the grassland associated fauna (Sukumar et al., 1995; Srinivasan, 2011; Vanak et al., 2013; Rawat and Adhikari, 2015). Therefore, it is important to incorporate these concerns while preparing the management plans for these grassland habitats.

The radio telemetry data from two female swamp deer indicate intensive use of the fragmented grassland patches outside protected area, corroborating with earlier information from Paul et al. (2018, 2020). While seasonal movements of swamp deer have been reported from the protected areas of central Indian landscape (Martin, 1977; Wildlife Institute of India, 2017) and Sharda river basin of north India (Ahmed, 2007) data from this study provided critical new insights on active migration routes, stopover sites and habitat use patterns in a highly human-dominated landscape. Analyses of the GPS locations identified non-protected areas between JJCR and Bijnor Barrage (Amichand, Nangal, Ranjeetpur, Sukhapur etc.) along the borders of Uttarakhand and Uttar Pradesh as highly used habitats (Supplementary Fig. A2 A). Some of these areas (Amichand and Sukhapur) are under severe pressure from land encroachment and overgrazing, and thus require immediate measures for protection of grasslands (Supplementary Fig. A2 E and F). BBMM analyses revealed grassland-dominated core home ranges of 0.58 (Female 1) and 1.47 (Female 2) km^2^ covering multiple grassland areas, suggesting importance of these fragmented patches. The habitat preference analyses (Ivlev’s index) also confirmed this pattern and showed high preferences towards grasslands. As availability of grassland is very low in landscape level compared to home ranges (landscape level- 6%; 95% BBMM- 40% and 24%; 50% BBMM- 79% and 46% for Female 1 and 2, respectively), the highest value of Ivlev’s index was observed at landscape scale (0.87 and 0.84, respectively for Female 1 and 2), suggesting obligatory grassland use by swamp deer. Apart from grasslands, cropland and waterbodies were also sporadically used by the animals (albeit with lower value than grasslands), possibly as they are the only transit routes available between the grassland patches. Further, fine-scale movement patterns (through trajectory path continuity analysis) support this as the results indicate that the swamp deer spend majority of the times within grassland patches and sporadically visit the croplands (probably for foraging or transit) and waterbodies (for transit). Croplands, in particular sugarcane fields also possibly provide covers during transit and facilitate movements but due to their seasonal availability (during monsoon) cannot serve as their prime habitats (Wurster et al., 2012; Athreya et al., 2013; Laforge et al., 2017). Even though temporal continuity within grasslands was highest for both individuals, fragmented grasslands with intersperse cropland and waterbodies may result in more temporal continuity within other classes (as observed in case of Female 2- 87% temporal continuity in cropland). Focused camera trapping exercise within the core habitats of the collared swamp deer (mainly outside protected areas) reveals small groups (4-5 individuals) inside the patches. This pattern supports known swamp deer ecology of summer congregations and seasonal movements in small groups in different landscapes (Martin, 1977; Ahmed, 2007). This strategy could allow them to maximise resource use and reduce confrontation with humans within the small grassland patches (Fahrig, 2007; Punjabi and Rao, 2017). Photographs of young fawns and rutting males within these core habitats also confirm that these patches are critical breeding and fawning grounds for the species (Supplementary Fig. A2 B and C). However, swamp deer showed temporal segregation with human, a behavioural strategy observed in other herbivores in human-dominated landscapes (Gaynor et al., 2018; Wilson et al., 2020; Naha et al., 2021). More radio-collaring and information from other areas in this landscape can substantiate the patterns found in this study.

Our analyses revealed that very small proportion (∼18%) of the study area (covering ∼53% of PA and ∼47% of non-PA) is suitable as swamp deer habitat. This specific region has already been identified as ‘Priority Conservation Areas’ in Paul et al. (2020), and our results substantiate the earlier selection. We believe that unlike the earlier study (Paul et al., 2020) availability of the accurate grassland positions (through surveys) and fine–scale location data (from both earlier study and radio collar GPS locations) has resulted in better understanding of the characteristics of these grassland patches. For example, quantification of the suitable habitats reveal that ∼69% (∼123 km^2^ out of ∼178 km^2^) of the grassland patches are identified as swamp deer suitable habitat, situated along the rivers Ganges and Solani. Further, majority of the highly suitable area (∼70% of 91 km^2^ area) are found outside the PA between JJCR and HWLS, supporting the earlier habitat use patterns from the collared animals. We found that most of the non-suitable grasslands are severely fragmented and away from the rivers, indicating the importance of the riverine system for swamp deer movement.

The vegetation mapping in this study is probably the first such information for this region. For vegetation mapping we selected four grassland species (*Saccharum* spp., *Phragmites* sp. and *Typha* sp.) in this work as earlier reports from Gangetic Plains and other areas suggested they are the major constituents of swamp deer habitats (Khan et al., 2003; Tewari and Rawat 2013b; Wildlife Institute of India, 2017). However, it is important to point out that other species of grasses such as *Cynodon* sp., *Imperata* sp. etc. are also reported from this landscape (Khan et al., 2003; Tewari and Rawat., 2013b). Our data from field surveys suggest that they are present sporadically in this landscape and possibly did not influence the classification schemes. The overall NDVI value can also get affected by the other noises such as presence of varying amount of water in the grassland patches, but we feel that it might be averaged out over reflectances from multiple patches. During the study period (April-May) *Saccharum* sp. was found to be the most dominant grass in the landscape, possibly due to their high colonisation ability during river course alteration events (Dinerstein, 2003; Johnsingh et al., 2004). Future efforts should focus to generate multi-season data on grass species to achieve perennial patterns of vegetation maps. Interestingly, we found that swamp deer (n=2 animals) mostly preferred (∼75%) Saccharum dominated mix and Phragmites dominated mix vegetations in this landscape during summer. This pattern is slightly different from earlier reports from JJCR (Tewari and Rawat, 2013b) where they preferred Typha over other vegetation types (Tewari and Rawat, 2013b). As availability of good habitat is limited across this human-dominated area, such preferences can vary based on local conditions (accessibility to vegetation type, disturbance regime, flooding patterns, seasonality etc.) and further detailed investigation is required to ascertain such patterns, if any.

## 5. Management recommendations

The results from this study lead to some suggestions regarding critical management/conservation interventions for the remaining grassland habitats in the upper Gangetic Plains. First, loss of most of the grassland habitats (∼57% reduction during last three decades) to croplands warrants immediate attention to the existing habitats in this landscape. As significant proportion of the most suitable habitat is situated at the border between Uttarakhand and Uttar Pradesh, a joint grassland management plan by both states is required for uniform implementation of any actions. Two particular interventions would be important: a) protection and recovery of the highly suitable areas of Amichand, Nangal, Ranjeetpur, Sukhapur etc. from encroachment and over-grazing as they contain large fragments of grasslands and form a ‘dynamic corridor’ between JJCR and HWLS to maintain a possible meta population framework for Gangetic fauna; and b) attention to appropriate management of the grassland habitats of HWLS. Despite being a protected area HWLS hosts a large number of human settlements and experiences various forms of anthropogenic pressure. Recent works have led to actions towards reappropriaton of HWLS boundary based on remaining habitats and biodiversity (Mondol et al., 2019) and future management efforts should ensure that after boundary realignment the habitat and biodiversity of HWLS is properly managed. This would require a collaborative approach with other government departments (Ministry of Agriculture and Farmers Welfare, Ministry of Housing and Urban Affairs, Department of Water Resources, River Development and Ganga Rejuvenation, Department of Revenue etc.) and proper plantation strategies, distribution of minimum numbers of agricultural licenses along rivers, review of the land tenure/revenue records, release of water from dams/barrages etc. should be properly planned.

During our field surveys we observed that the grasslands habitats contain mixture of swampy tall grassland/wetland (dominated by *Phragmites* sp., *Typha* sp. and *Saccharum* spp. with perennial water availability) and dry grasslands (dominated with *Saccharum* spp. mostly along riverbeds) (Johnsingh et al., 2004; Rawat and Adhikari, 2015). These grassland habitats require different management strategies involving appropriate burning regimes, water level control etc. for their long-term survival. Their persistence will also ensure viability of their respective faunal biodiversity (fishing cat (*Prionailurus viverrinus*), hog deer (Axis porcinus), freshwater crocodile (*Crocodylus johnsoni*), gharial (*Gavialis gangeticus*), painted stork (*Mycteria leucocephala*), Indian saras crane (*Antigone Antigone*) in wet grasslands and Bengal florican (*Houbaropsis bengalensis*), hispid hare (*Caprolagus hispidus*)and blackbuck (*Antilope cervicapra*) in dry grasslands (Odden et al., 2005; Varghese et al., 2008; Mukherjee et al., 2012; Nath and Machary, 2015; Thakuri, 2018; BirdLife International, 2021).

## 6. Conclusion

In conclusion, our findings in this study provide a fine-scale baseline information of the distribution, dynamics and vegetation maps of the high faunal biodiversity retaining grasslands in the upper Gangetic Plains. Recent reports suggest that with a loss of ∼20 million hectares area during last century the grassland ecosystems are one of the most neglected in India (Report of the task force on Grasslands and Deserts, 2006). Global projection of biodiversity change scenarios identified grasslands as one of the most vulnerable ecosystems in coming century (Sala et al., 2000, 2005). We believe that the quantitative information provided in this study would be considered as a model to generate more information and help restoring such rapidly vanishing ecosystem worldwide.

## 7. Acknowledgement

We acknowledge the Forest Departments of Uttarakhand, Uttar Pradesh and MOEFCC (No. 244/2018/RE), Government of India for providing necessary permits to carry out the research. Our thanks to the Forest Department officials especially Divisional Forest Officer, Haridwar Forest Division and local community members for their assistance in field work. We acknowledge help from Vinod Thakur, Tista Ghosh, Suvankar Biswas, Dr Sarbesh Rai, Shiv Kumari Patel, Shrushti Modi, Kaushal Singh Sissodia, Sultan Singh Badana, Nimisha Srivastava, Rakesh Mondol, Debanjan Sarkar, Prajak Kumar Das, Chitrapal, Imam, Ranju, Bhura, Annu, Juri, Inam, Ammi, Msc students of Wildlife Institute of India (XVI batch) and villagers of Gendikhata and Tantwala for their help during the collaring operation and field surveys. We appreciate technical help from Dr Gautam Talukder and Aishwariya Ramachandran for GIS work. We thank the Director, Dean and Research Coordinator of Wildlife Institute of India for their support. The work was funded by Uttarakhand Forest Department, Uttar Pradesh Forest Department, MOEFCC, Governnmnet of India and Conservation Force. Shrutarshi Paul was awarded Department of Science and Technology INSPIRE Research Fellowship (IF150680) and Samrat Mondol was supported by Department of Science and Technology INSPIRE Faculty Award (IFA12-LSBM-47).

## CRediT authorship contribution statement

**Shrutarshi Paul:** Conceptualization, data curation, formal analysis, methodology, writing-original draft, writing review and editing

**Sohini Saha:** Data curation, methodology, formal analysis, writing-review and editing

**SK Zeeshan Ali:** Methodology, resources, writing- review and editing

**Navendu Page:** Methodology, supervision, writing- review and editing

**Parag Nigam:** Methodology, supervision, resources, writing- review and editing

**Aamer Sohel Khan:** Data curation, formal analysis, writing-review and editing

**Mukesh Kumar:** Conceptualization, supervision, resources, writing- review and editing

**Bilal Habib:** Methodology, resources, writing- review and editing

**Dhananjai Mohan:** Conceptualization, supervision, writing- review and editing

**Bivash Pandav:** Conceptualization, funding acquisition, project administration, supervision, resources, writing- review and editing

**Samrat Mondol:** Conceptualization, funding acquisition, project administration, supervision, resources, investigation, writing-original draft, writing review and editing

## Figure legends

**Supplementary Fig. A1:**
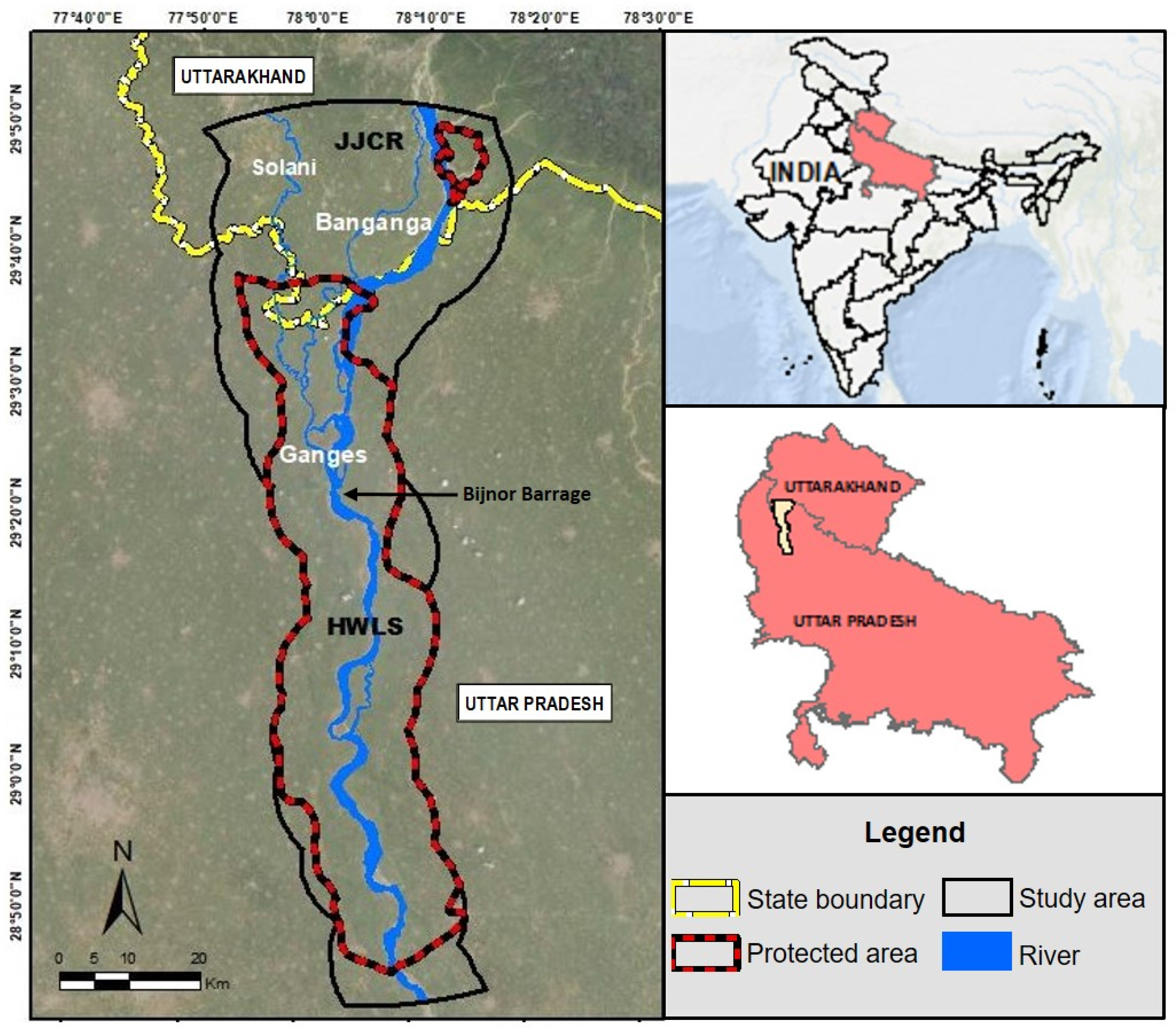
Map of the study area covering the states of Uttarakhand and Uttar Pradesh. The study area consists of two protected aeas (JJCR and HWLS) and other non-protected regions.

**Supplementary Fig. A2:**
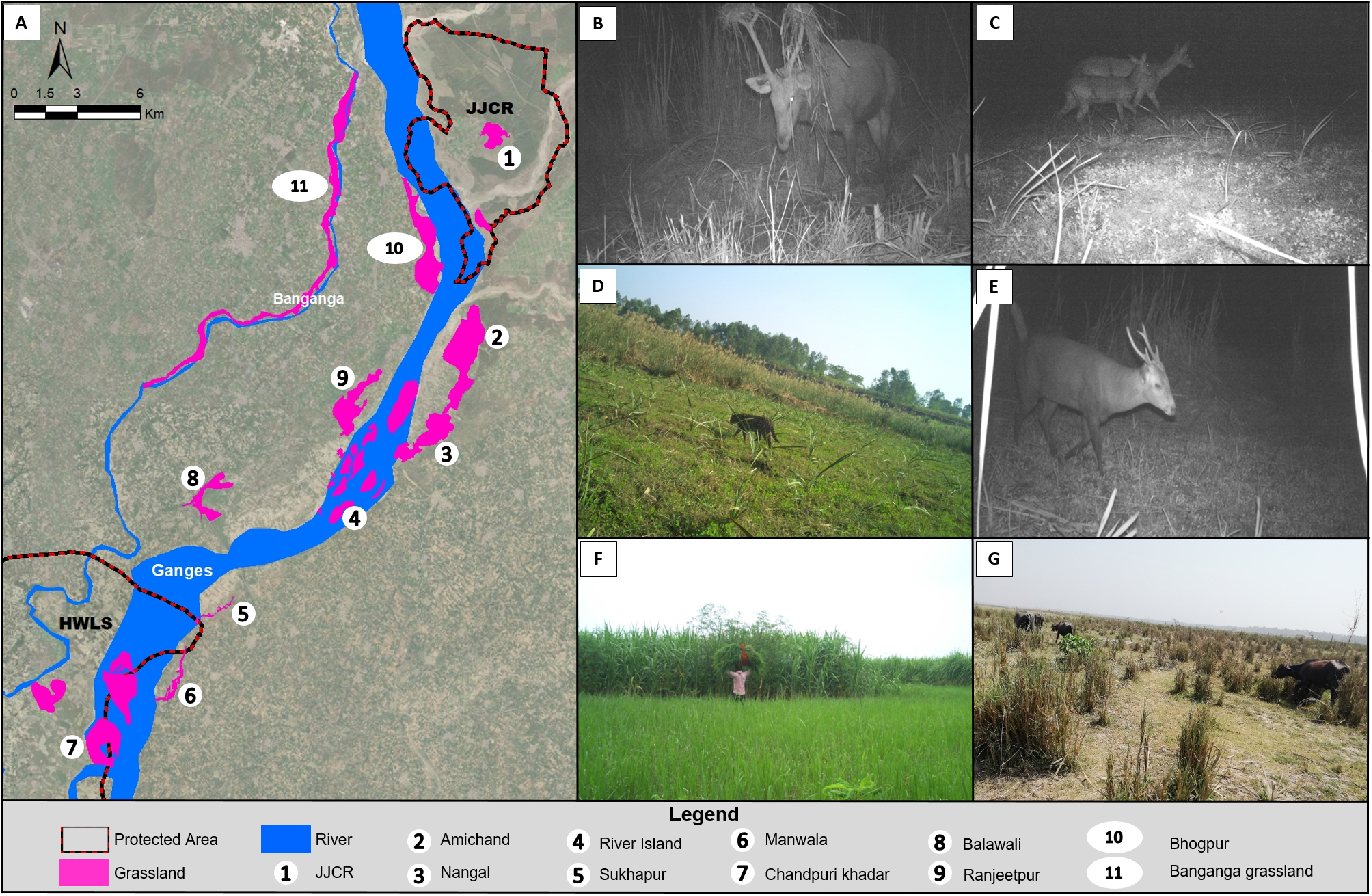
Grassland habitats, biodiversity and disturbances encountered in some of non-protected areas between JJCR and HWLS (A) Digitized locations of major grassland patchs between the two protected areas. Based on the habitat use patterns from the collared females, camera trapping was conducted in locations 2, 3, 4, 6, and 9; (B) Camera trap image showing rutting behaviour of a male during late monsoon; (C) Capture of swamp deer fawn with a female indicating the importance of the grassland habitats as fawning and breeding grounds; (D) Evidence of fishing cat using the habitats; (E) Evidence of hog deer in these habitats; (F) Human disturbances in the form of exploitation/extraction of grasses; (G) Overgrazing of the habitats by live stock.

**Supplementary Fig. A3:**
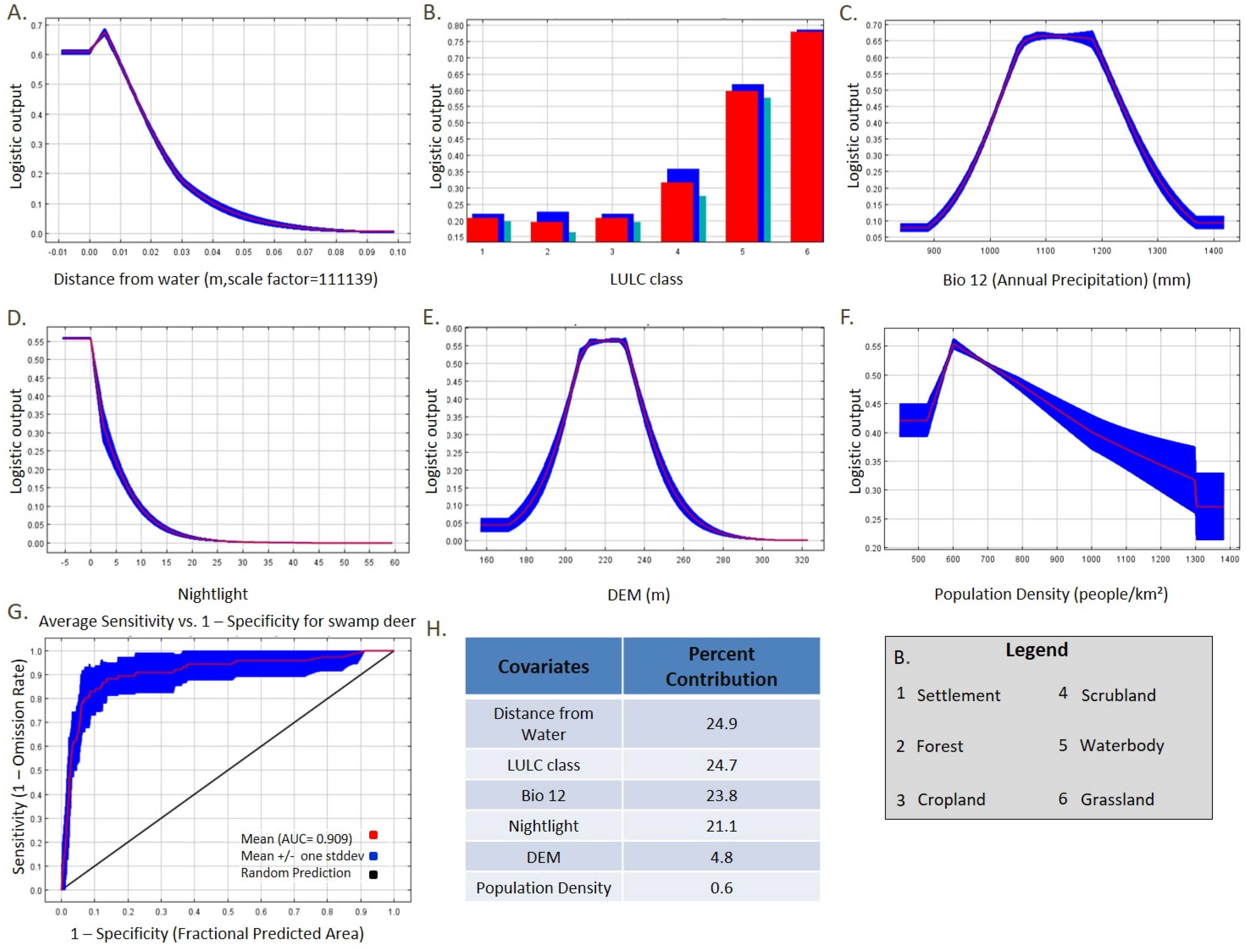
Response curves of six covariates used in MaxEnt analyses to ascertain swamp deer habitat suitability. Figures A-F shows respective response curves and G lists the estimates of relative contributions of these covariates in the MaxEnt predictions. Figure H shows the ROC curve for the training omission rate and predicted area averaged over the replicate runs.

**Supplementary Fig. A4:**
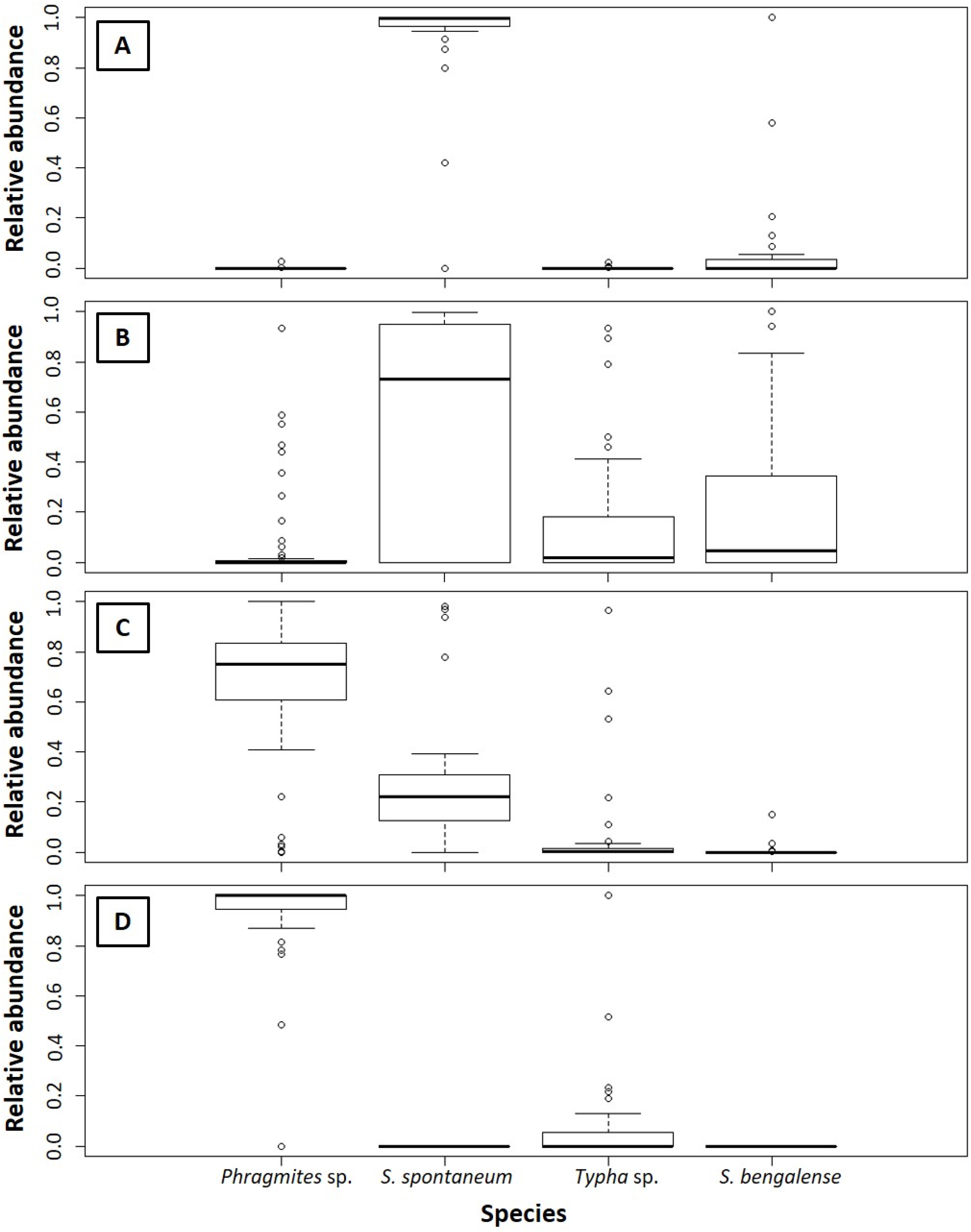
Relative abundances of four grass species (*Phragmites* sp, *S. spotaneum*, *Typha* sp and *S. bengalense*) in each vegetation types:(A) pure Saccharum; (B) Saccharum Dominated Mix; (C) Phragmites Dominated Mix; (D) pure Phragmites/ Typha

**Supplementary Fig. A5:**
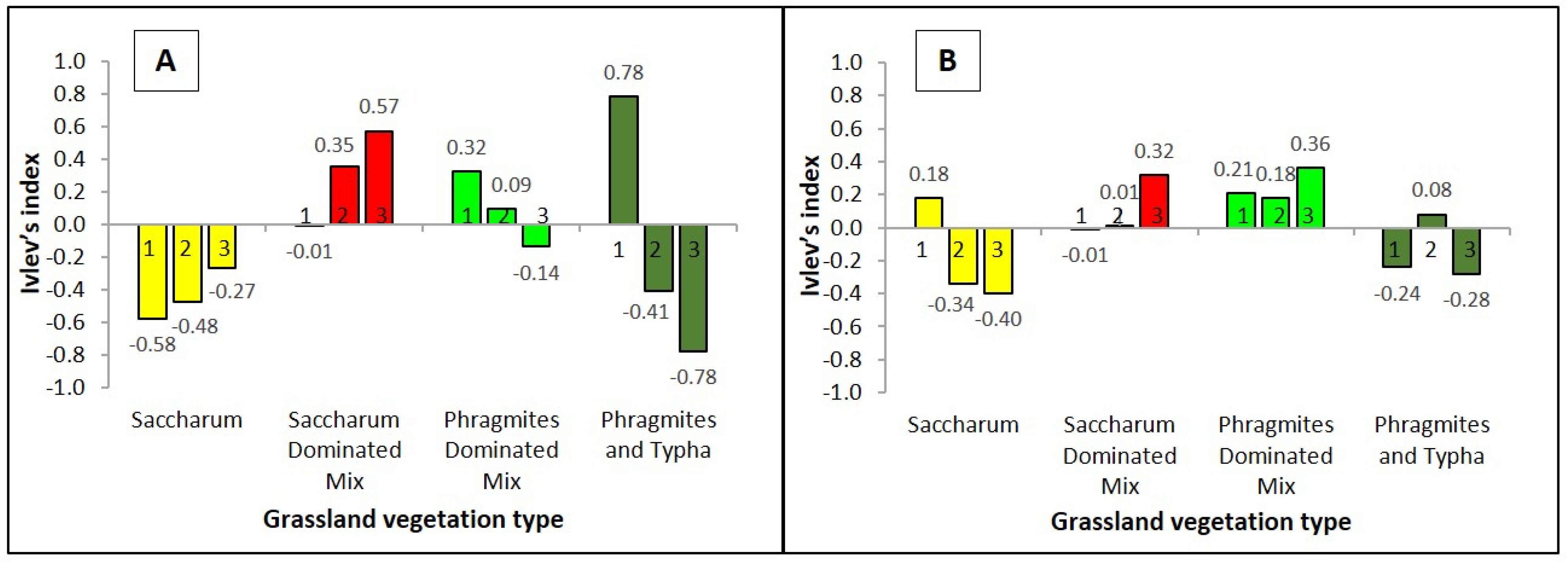
Preferences towards different vegetation types (Ivlev’s Index) within grassland for two collared females (represented in A and B) during summer. In all cases, 1 depicts 50% BBMM, whereas 2 and 3 represents 95% BBMM and the entire landscape.

**Supplementary Fig. A6:**
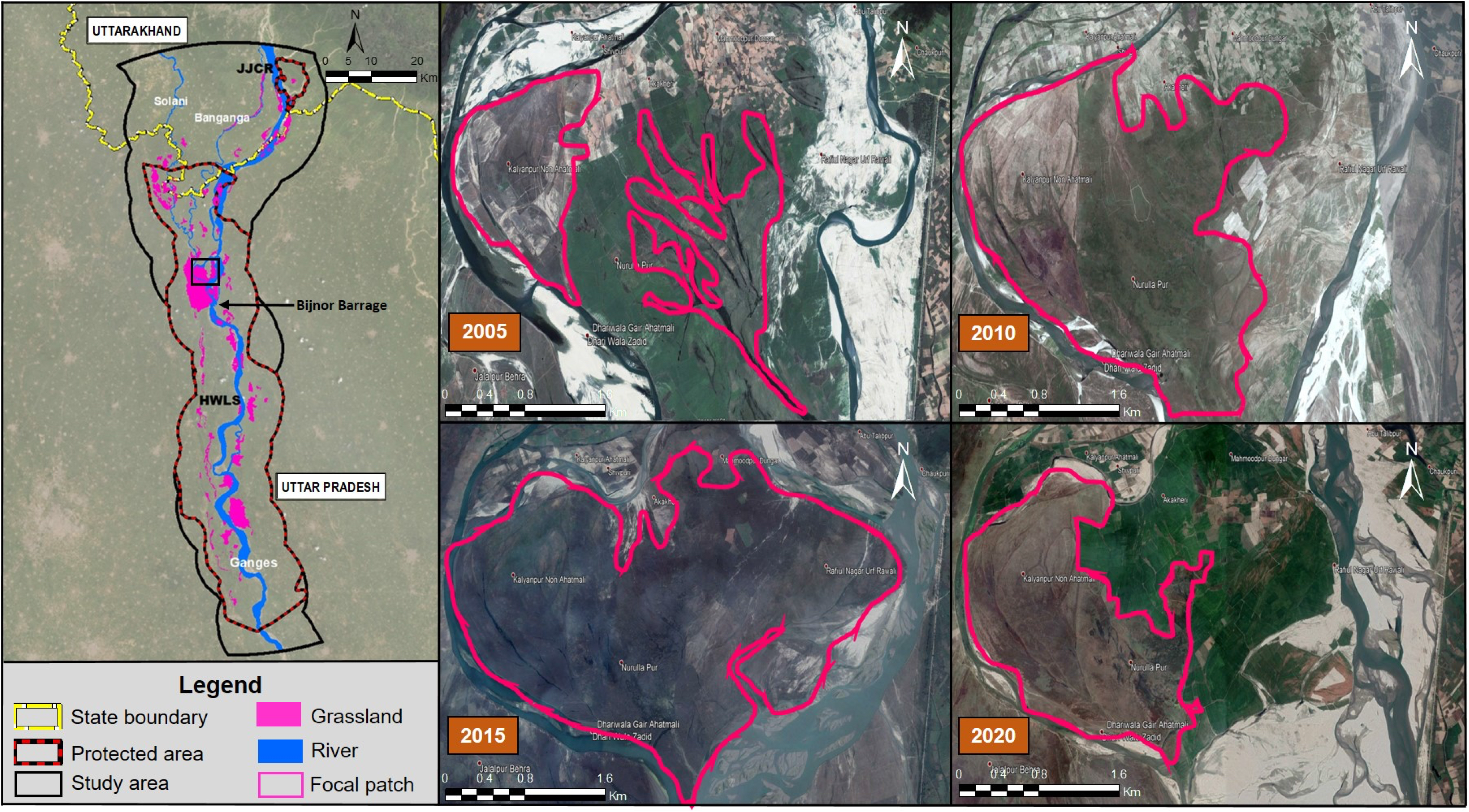
Times-series presentation of grassland dynamics within a selected river island area (Rauli Ghat, presented in the left pane of the figure) of the upper Gangetic Plains. The changes in the grassland habitat area are presented at every 5-year time frame during last two decades (2005, 2010, 2015 and 2020).

## Table legends

**Supplementary Table A1:** Mean relative abundance of *Phragmites* sp., *Saccharum spontaneum*, *Typha* sp. and *Saccharum bengalense* within the four vegetation classes

| Vegetation Class | Mean relative abundance | | | | Kruskal Wallis ($\chi^2$) |
| --- | --- | --- | --- | --- | --- |
|  | *Phragmites* sp. | *Saccharum spontaneum* | *Typha* sp. | *Saccharum bengalense* |  |
| **Saccharum** | < 0.001 | 0.85 | <0.001 | 0.14 | 96.60* |
| **Saccharum Dominated Mix** | 0.06 | 0.57 | 0.12 | 0.23 | 48.48* |
| **Phragmites Dominated Mix** | 0.66 | 0.26 | 0.07 | <0.001 | 96.35* |
| **Phragmites and Typha** | 0.87 | 0 | 0.13 | 0 | 136.23* |

\*p<0.05

